# The Ras-like GTPase Rem2 is a potent endogenous inhibitor of calcium/calmodulin-dependent kinase II activity

**DOI:** 10.1101/148981

**Authors:** Leandro Royer, Josiah J. Herzog, Katelyn Kenny, Boriana Tzvetkova, Jesse C. Cochrane, Michael T. Marr, Suzanne Paradis

**Affiliations:** Department of Biology, Brandeis University, Waltham, Massachusetts 02454; Department of Molecular Biology and Genetics, Massachusetts General Hospital and Harvard Medical School, Boston, Massachusetts 02114; Rosenstiel Basic Medical Sciences Research Center; Volen Center for Complex Systems; National Center for Behavioral Genomics, Brandeis University, Waltham, Massachusetts 02454

**Keywords:** calcium, Ca2+/calmodulin-dependent protein kinase II, GTPase, inhibitor, phosphorylation, Rem2, RGK family

## Abstract

CaMKII is a well-characterized, abundant protein kinase that regulates a diverse set of functions in a tissue specific manner. For example, in heart muscle, CaMKII regulates Ca^2+^ homeostasis while in neurons CaMKII regulates activity-dependent dendritic remodeling and Long Term Potentiation (LTP), a biological correlate of learning and memory. Previously, we identified the noncanonical GTPase Rem2 as a critical regulator of dendrite branching and synapse formation in the vertebrate nervous system. Here, we report that Rem2 directly interacts with CaMKII and potently inhibits the activity of the intact holoenzyme, a previously undescribed function for the Rem2 protein. To date, only one other endogenous inhibitor of CaMKII has been described: CaMKIIN, which blocks CaMKII activity through binding to the catalytic domain. Our data suggest that Rem2 inhibits CaMKII through a novel mechanism, as inhibition requires the presence of the association domain of CaMKII. Our biochemical finding that Rem2 is a direct, endogenous inhibitor of CaMKII activity, coupled with known functions of Rem2 in neurons, provides a framework which will enable future experiments probing the physiological role of CaMKII inhibition in a cellular context.

## Introduction

Calcium/calmodulin-dependent kinase II (CaMKII)^2^ is an abundant, multifunctional serine-threonine kinase whose regulation and activity is uniquely sensitive to intracellular Ca^2+^ levels (1-3). Modulation of CaMKII by Ca^2+^ is one of its most interesting features: Ca^2+^/calmodulin binds to the regulatory region of CaMKII leading to exposure of the catalytic domain and activation of the enzyme (4). Subsequent phosphorylation at Thr286 on the regulatory domain by an adjacent CaMKII monomer in the holoenzyme leads to an autonomous, Ca^2+^-independent kinase activity. Importantly, autonomous CaMKII is temporally uncoupled from the original cellular signal that led to its activation, potentially allowing the enzyme to act as a molecular memory of previous Ca^2+^ signaling events (5,6).

The functions of CaMKII are best understood in excitable cell types such as neurons and heart muscle, where Ca^2+^ signaling plays a critical role in the functional output of these cells. The unique properties of CaMKII activation allow the enzyme to integrate both the frequency and amplitude of Ca^2+^ signaling, and translate this information into specific cellular outcomes (7-9). Well-studied substrates of CaMKII include ion channels such as AMPA receptors, voltage-gated calcium and potassium channels, and the transcription factor CREB (10-12). In the nervous system, activation of CaMKII by Ca^2+^ entry during high-frequency stimulus plays a crucial role in synaptic plasticity induction and maintenance (1,9,13). This fact, in conjunction with CaMKII’s autonomous function, led to the hypothesis that this kinase is a major molecular determinant of learning and memory (14). In heart muscle CaMKII is a major actor in Ca^2+^ homeostasis, regulating the cardiomyocyte excitability and contractility through the modulation of several channels including the L-type voltage-gated Ca^2+^ channel (15,16). Accordingly, recent studies determined that CaMKII deregulation is an important element in the pathogenesis of arrhythmias and heart failure (17-21), with CaMKII inhibition suggested as a potential therapy. Despite intense research in several fields for over thirty years, we still do not completely understand how the activation status of CaMKII is regulated. Thus, there is intense interest in achieving a better understanding of CaMKII regulation both from a basic science and human health perspective.

The kinase activity of CaMKII is regulated in part by cytosolic phosphatases (22). However, during certain cellular processes such as synaptic potentiation, CaMKII accumulates in cellular regions with low local concentration of phosphatases (23,24). How uncontrolled kinase activity is avoided in those circumstances is still not completely understood. One potential mechanism would be the activity-regulated expression of endogenous CaMKII inhibitors. Nevertheless, despite intensive research into CaMKII regulation, molecular mechanisms of CaMKII inhibition are not well understood. In fact, only one endogenous inhibitor has been described in mammals (25,26). Here we present data demonstrating that the small GTPase Rem2 interacts with CaMKII and is a novel, endogenous inhibitor of CaMKII activity.

Rem2 is a member of the RGK sub-family (Rem2, Rad, Rem and Gem/Kir) of the Ras superfamily of monomeric G-proteins (27). A number of features differentiate RGK family members from the Ras superfamily. For example, the crystal structures of several RGK proteins, including Rem2, reveal differences between this family and classical GTPases in the structure of their nucleotide binding domains (28,29). For the most part, neither GTPase activating proteins (GAPs) nor guanine nucleotide exchange factors (GEFs) have been identified for this family (30) (the only exception being the identification of NmeI as a Rad GAP (31)). RGK proteins contain extended N and C-termini relative to other Ras family members (30), but lack the C-terminal CAAX domain, which serves as a prenylation signal and targeting to the plasma membrane (32). For these and other reasons, Rem2 and the RGK family members in general may not behave as classical Ras-like GTPases regulated by their nucleotide binding state.

Overexpression of RGK family members in various cell types has been shown to profoundly inhibit high voltage-activated Ca^2+^ channels (30,33-35). The functional consequences of the decreased Ca^2+^ influx through voltage-gated calcium channels caused by RGK proteins is not completely characterized, nor is the mechanism by which RGK proteins mediate this effect. We have previously shown that Rem2 mRNA expression is upregulated by neuronal activity (36). Moreover, using gene knockdown approaches in cultured cortical neurons and *X. laevis* optic tectum we have established key functions of Rem2 in the nervous system: Rem2 acts as a positive regulator of synapse formation and a negative regulator of dendritic branching (36,37). As an inroad to obtaining a detailed understanding of Rem2 signaling we took an unbiased approach to identifying Rem2 interacting proteins and found that Rem2 directly interacts with CaMKII. Further, using *in vitro* kinase assays, we demonstrate that Rem2 inhibits the activity of CaMKII, and that inhibition requires the CaMKII association domain, suggesting a unique mechanism. Our finding that Rem2 is a direct, endogenous inhibitor of CaMKII activity has important implications for regulation of synaptic plasticity and activity-dependent neuronal remodeling in the mammalian brain.

## Results and Discussion

To identify molecules that potentially interact with Rem2 in cells, we performed a biochemical enrichment of Rem2-interacting proteins followed by mass spectrometry identification. We passed a whole brain lysate from P14 rats over a column containing a Maltose Binding Protein (MBP)- Rem2 fusion protein bound to amylose-agarose or a control MBP alone column. We identified specifically bound proteins by mass spectrometry. Using this approach, we found that all four isozymes of CaMKII interact with Rem2 (Fig. 1A). Although Rem2 is a substrate of CaMKII *in vitro* (38) our ability to detect the CaMKII-Rem2 interaction by affinity chromatography suggests a more stable interaction between Rem2 and CaMKII than that which is typical between a kinase and substrate. Consistent with this, Rem2 has been previously shown to interact with CaMKIIα (39). We went on to confirm this interaction by co-immunoprecipitating an HA epitope-tagged Rem2 protein (HA-Rem2) and a myc epitope-tagged CaMKIIα protein expressed in HEK 293T cells using an anti-HA antibody (Fig. 1B).

**Figure 1.**
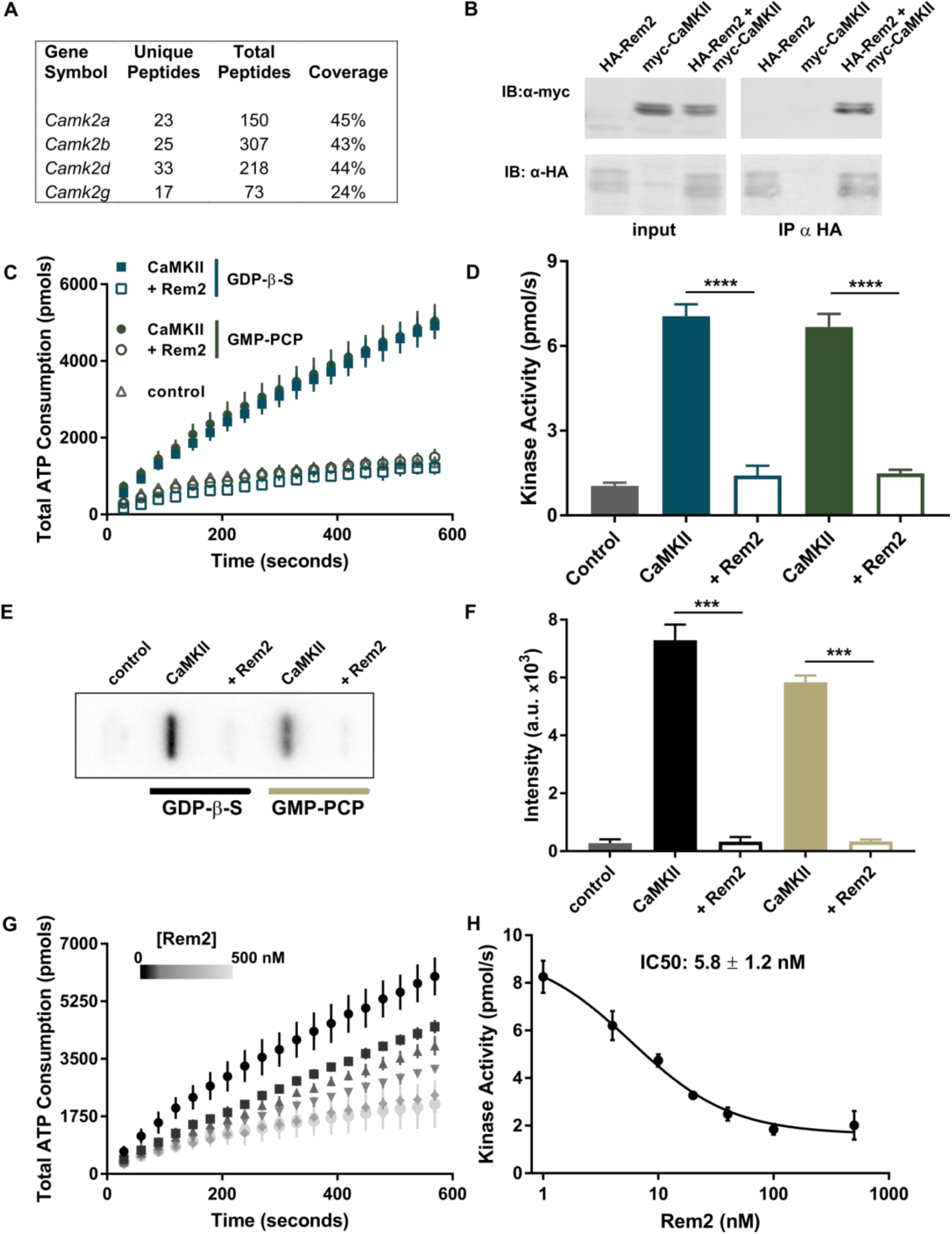
Rem2 binds to CaMKII and inhibits its kinase activity. (A) Mass spectrometry data identifies CaMKII isozymes retained by a Rem2 affinity matrix after loading with total brain lysate from P14 wild-type rats. The number of unique and total peptides and the percentage of the protein sequence covered for each isoform is shown. (B) Co-immunoprecipitation of Rem2 and CaMKII from HEK293T cells transfected with HA-tagged Rem2, myc-tagged CaMKIIα or both. Rem2 was immunoprecipitated using an anti-HA antibody and anti-myc and anti-HA immunobloting was performed to detect CaMKII and Rem2, respectively. (C) ATP consumption by purified CaMKIIα (20 nM monomer) monitored using the PK/LDH assay with syntide-2 as a substrate (200 μM) with 500 nM Rem2 (filled symbols) or without 500 nM Rem2 (hollow symbols) in the presence (blue squares) of 40 μM GDP-β-S or 40 μM GMP-PCP (green circles). Samples devoid of CaMKIIα were used as negative controls (gray triangle). (D) Quantification (see methods) of CaMKIIα kinase activity shown in (C) (N = 6 reactions; bars indicate standard deviations; ****, p < 0.0001, using Student’s t-test – see Materials and Methods for details on the statistical analysis). (E) Representative autoradiograph of a slot blot of ^32^P-labeled syntide-2. The phosphorylation of syntide-2 (200 μM) by CaMKIIα (20 nM) was conducted in the presence of radiolabeled ATP for 8 min in the presence (500 nM) or absence of Rem2 in the presence or either 40 μM GDP-β-S or 40 μM GMP-PCP as indicated. Sample without syntide-2 was used as a control. (F) Quantification of all syntide-2 autoradiographies as in panel (E) by densitometry of the bands. (N = 4; standard deviations shown; ***, p < 0.001, using Student’s t-test). (G) Concentration-dependence of Rem2 inhibition of CaMKIIα kinase activity towards syntide-2. Black circles 0 nM Rem2; black squares, 4 nM Rem2; upward triangles 10 nM Rem2; downward triangles 20 nM Rem2; gray diamonds 40 nM Rem2; light grey circles 100 nM Rem2. (H) Dose-response plot of the data shown in panel (G). The average value of the slope of each curve was plotted as a function of Rem2 concentration (anti-log scale). The half-maximal inhibitory concentration (IC50) is found at 5.8 ± 1.2 nM and was obtained from the non-linear fit of the points (see Materials and Methods).

To examine the relationship between Rem2 and CaMKII we took an *in vitro* approach to ask what effect Rem2 has on CaMKII activity. We purified recombinant Rem2 and CaMKIIα holoenzyme, the main isozyme expressed in the adult brain (40), from *E. coli*. Negative stained EM images of the peak fraction confirmed that the purified CaMKIIα is present as the holoenzyme dodecamer (Fig. S1). To test the kinase activity of the enzyme we employed the classic Pyruvate Kinase/Lactate Dehydrogenase coupled assay (PK/LDH) (9,41). In this assay, the production of ADP by the phosphorylation reaction of the kinase is coupled to the decay of NADH to NAD^+^ in a one to one molar ratio. Thus, monitoring the rate of NADH decay by spectrophotometry provides a quantification of ADP production and thus ATPase activity. Using this assay, we determined that our preparation of CaMKIIα is active using syntide-2 as a substrate (Fig. 1C-D).

Interestingly, CaMKIIα is strongly inhibited by addition of Rem2 to the assay (Fig. 1C-D, Fig. S2). In fact, Rem2 completely inhibits CaMKIIα, limiting ADP production to a level equal to our control condition that lacks CaMKIIα. Using non-hydrolyzable analogs of GTP (GMP-PCP) or GDP (GDP-β-S) we found no difference in the ability of Rem2 to inhibit CaMKII (Fig. 1C-D). It is important to note that under our experimental conditions, the concentration of guanine nucleotides used (40 μM) is saturating for Rem2 but has been shown to not affect the activity of CaMKIIα (27,42). Accordingly, addition of the nucleotides in the absence of Rem2 did not alter the kinase activity of CaMKII (Fig. 1C-D). This result suggests that the mechanism of inhibition is independent of which nucleotide is bound to Rem2. Members of the RGK family have been shown to be non-canonical G-proteins with respect to the regulatory role of their G-domain (43). To confirm this result using an independent method, we performed a kinase assay using [γ-^32^P]ATP under the same conditions. In agreement with our observations using the PK/LDH assay, we found that phosphorylation of the syntide-2 substrate by CaMKIIα is significantly decreased in the presence of Rem2 (Fig. 1E-F). In addition, titration of Rem2 protein in our PK/LDH assay indicates that Rem2 is a potent inhibitor of CaMKII with an apparent IC50 of ~6 nM using the peptide syntide-2 as substrate (Fig. 1G-H). For comparison, the IC50 on the same substrate of the only other mammalian, endogenous CaMKII inhibitor, CaMKIIN, is 50 nM (26).

CaMKII inhibition could occur via a number of possible mechanisms, which would have different functional consequences for CaMKII signaling. For example, an inhibitor could act on the activation step, block the autophosphorylation of CaMKII, or block the catalytic activity. In the first case, the block of enzyme activation would result in silencing all signaling events mediated by CaMKII. Alternatively, interfering with the autophosphorylation of CaMKII would allow the enzyme to be active only when Ca^2+^/CaM is bound, effectively converting CaMKII into a signaling component without autonomous activity, preventing the creation of molecular memory. Finally, an inhibitor of the catalytic activity against exogenous substrates would allow the autophosphorylation of CaMKII but would prevent phosphorylation of other targets. This type of inhibition could be relevant in repetitive stimulation, permitting the formation of molecular memory but avoiding excessive phosphorylation of protein targets. Thus, in order to clarify the potential functional consequences of CaMKII inhibition by Rem2 we began an investigation of the properties of this inhibition.

First, we sought to determine if Rem2 interferes with the activation of CaMKII by Ca^2+^/CaM. We titrated the amount of CaM present in the PK/LDH assay in the presence or absence of 500 nM Rem2 (Fig. 2A-B, Fig. S3A). We found that Rem2 efficiently inhibited CaMKII activity over a wide range of CaM concentrations equal to or above the Rem2 concentration in the assay (i.e., 40-fold, 0.5- 20 μM), indicating that Rem2 is unlikely to act simply as a CaM scavenger. Next, we asked if Rem2 inhibited CaMKII autophosphorylation. In this experiment, autophosphorylation of CaMKIIα was induced by incubation of the enzyme with Ca^2+^/CaM and [γ-^32^P]ATP in the presence or absence of 500 nM Rem2 for 2 min. This incubation time has been shown to produce strong phosphorylation at the CaMKII autophosphorylation site residue Thr286 (44). The reaction mixture was separated by SDS-PAGE followed by autoradiography of ^32^P-labeled CaMKIIα (Fig. 2C-D). We found no difference in CaMKII autophosphorylation in the presence or absence of Rem2. This result indicates that Rem2 does not inhibit CaMKII autophosphorylation suggesting that it is very unlikely that Rem2 acts by hindering CaM binding to the regulatory domain of CaMKII. In addition, we observed phosphorylation of Rem2 by CaMKII as expected (38). Taken together, these experiments suggest that Rem2 acts as an inhibitor of CaMKII catalytic activity onto substrates other than the CaMKII regulatory domain.

**Figure 2.**
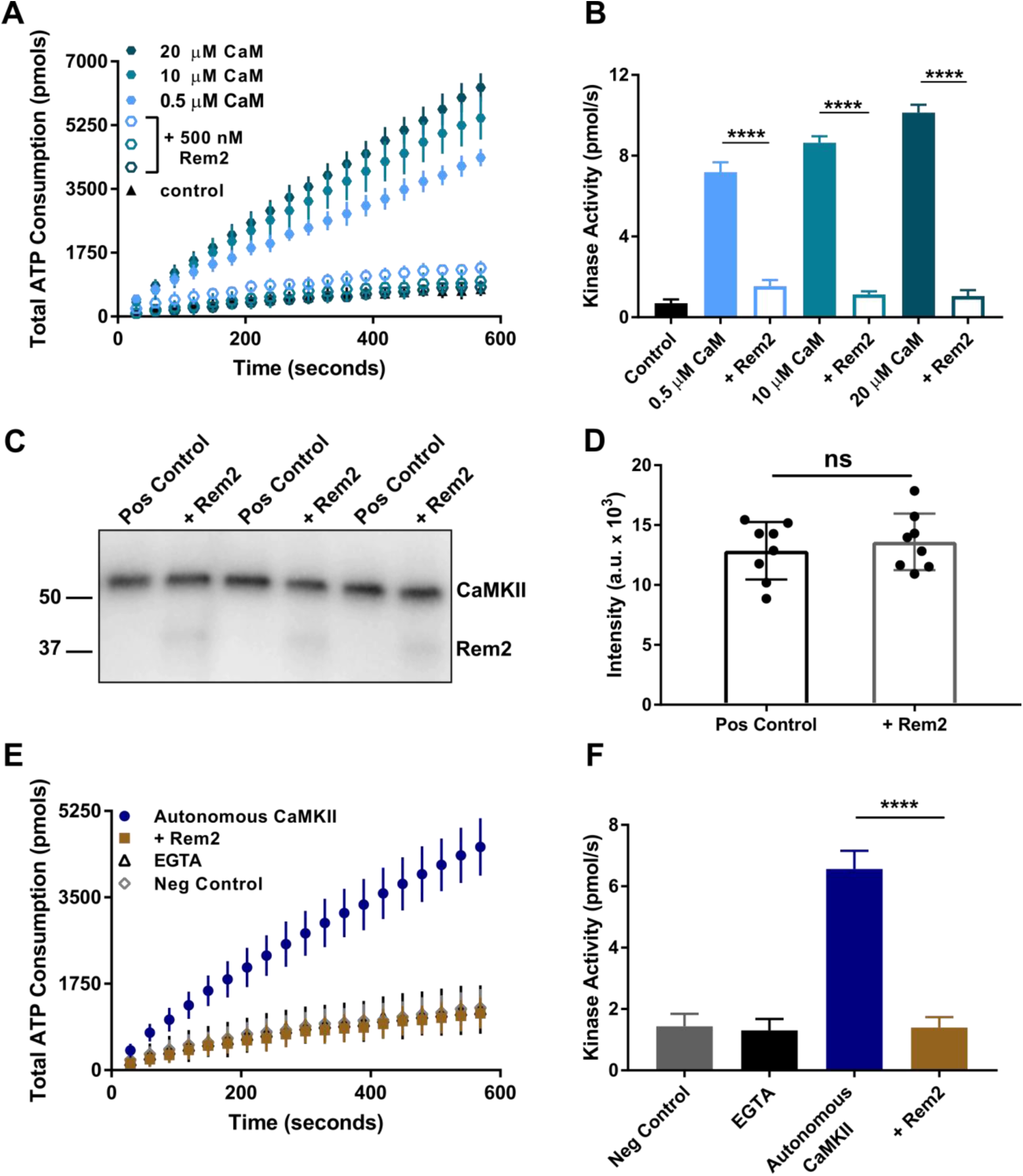
Rem2 inhibits autonomous CaMKIIα activity but not its autophosphorylation. (A) Total ATP consumption during the phosphorylation of syntide-2 (200 μM) by α (20 nM) was monitored at different concentrations of CaM (filled circles; 40-fold range). Addition of 500 nM Rem2 (open circles) reduced the ATP consumption to control levels (triangles) at any CaM concentration tested. (B) Quantification of the CaMKIIα kinase activity shown in (A) using the same color code (N = 6 for all CaM only conditions; N = 4 for 0.5 μM + Rem2 condition; N = 5 for all other conditions; error bars depict standard deviations; ****, p < 0.0001, using Student’s t-test). (C) Autophosphorylation of CaMKIIα (80 nM) was induced by incubation of the enzyme with Ca^2+^/ CaM (1 mM/3.33 μM) and [γ-^32^P]ATP (100 μM) in the presence or absence of 500 nM Rem2 for 2 min. The reaction mix was separated by SDS-PAGE, the gel was dried and visualization of ^32^P-labeled proteins was obtained using a phosphorimager. One representative experiment is shown; experiment was performed in triplicate. The numbers represent the position of protein molecular weight standards in kDa. (D) Quantification of all CaMKIIα autophosphorylation experiments as in (C) by densitometry (N = 8; standard deviations shown; ns, not significantly different (p = 0.5440), using Student’s t-test). (E) CaMKIIα (60 nM) was pre-incubated with ATP (500 μM) and Ca^2+^/ CaM (1 mM/10 μM) for 2 min at room temperature to promote CaMKIIα autophosphorylation and CaMKIIα autonomous activity. The pre-incubated mix was then exposed to syntide-2 (500 μM) ([Ca^2+^] < 50 nM). The total ATP consumption as a function of time in the absence (blue circles) or presence of Rem2 (500 nM, gold squares) is shown. Addition of EGTA during the pre-incubation was used to confirm that only the autonomous activity was assayed (black triangles). A negative control (gray diamonds) was conducted in the absence of CaMKIIα. (F) The kinase activity of CaMKIIα in each of the conditions described in panel (E) is shown. The same color code used in (E) is applied (N = 6; error bars depict standard deviations; ****, p < 0.0001, using Student’s t-test).

To test this hypothesis, we sought to determine if Rem2 interferes with CaMKII catalytic activity on syntide-2 by asking if Rem2 inhibits the Ca^2+^/CaM-independent, autonomous form of the enzyme. CaMKIIα was pre-incubated with ATP and Ca^2+^/CaM for 2 min to promote CaMKIIα autophosphorylation at residue Thr286 and CaMKIIα autonomous activity (44). After the incubation, free calcium in the solution was brought to values below 50 nM by addition of EGTA to suppress any Ca^2+^/CaM dependent activity. The pre-incubated mix was then immediately used in the PK/LDH assay in the presence or absence of 500 nM Rem2 using syntide-2 as a substrate (Fig. 2E-F, Fig. S3B). As a negative control, we added EGTA during the CaMKII pre-incubation to confirm that only the CaMKII autonomous activity was assayed (Fig. 2E, black triangles). We found that addition of Rem2 after CaMKII pre-incubation caused a severe decrease in total ATP consumption, indicating a strong inhibition of the autonomous activity of CaMKIIα.

All CaMKII substrates bind to the S-site on the catalytic domain of the enzyme. However, a subset of the substrates also bind to the Thr286- binding site on the catalytic domain, known as the T-site (45). Thus, the two types of substrates, S-type and T-type, have different interactions with the catalytic domain of CaMKII and selective modulation of the function of these sites could potentially provide an additional layer of regulation on CaMKII signaling. In neurons, the two classes of ionotropic glutamate receptors involved in synaptic plasticity, AMPA receptors and NMDA receptors, are examples of S- and T-type substrates, respectively (45). Since we observed that Rem2 inhibits the catalytic activity of CaMKII, we next asked whether Rem2 selectively inhibits CaMKII phosphorylation of specific types of substrates. Using the PK/LDH assay we compared the efficacy of Rem2 inhibition of CaMKII using the peptides syntide-2 or autocamtide-2 as models of S- and T-type substrates, respectively (Fig. 3A-B, Fig. S4A). We found that Rem2 is a much more potent inhibitor of CaMKII activity on syntide-2, indicating a preference for Rem2 inhibition of S-type substrates, thus hinting at a substrate specific role for Rem2 inhibition. A similar selectivity has been described for the CN21a inhibitory peptide (46).

**Figure 3.**
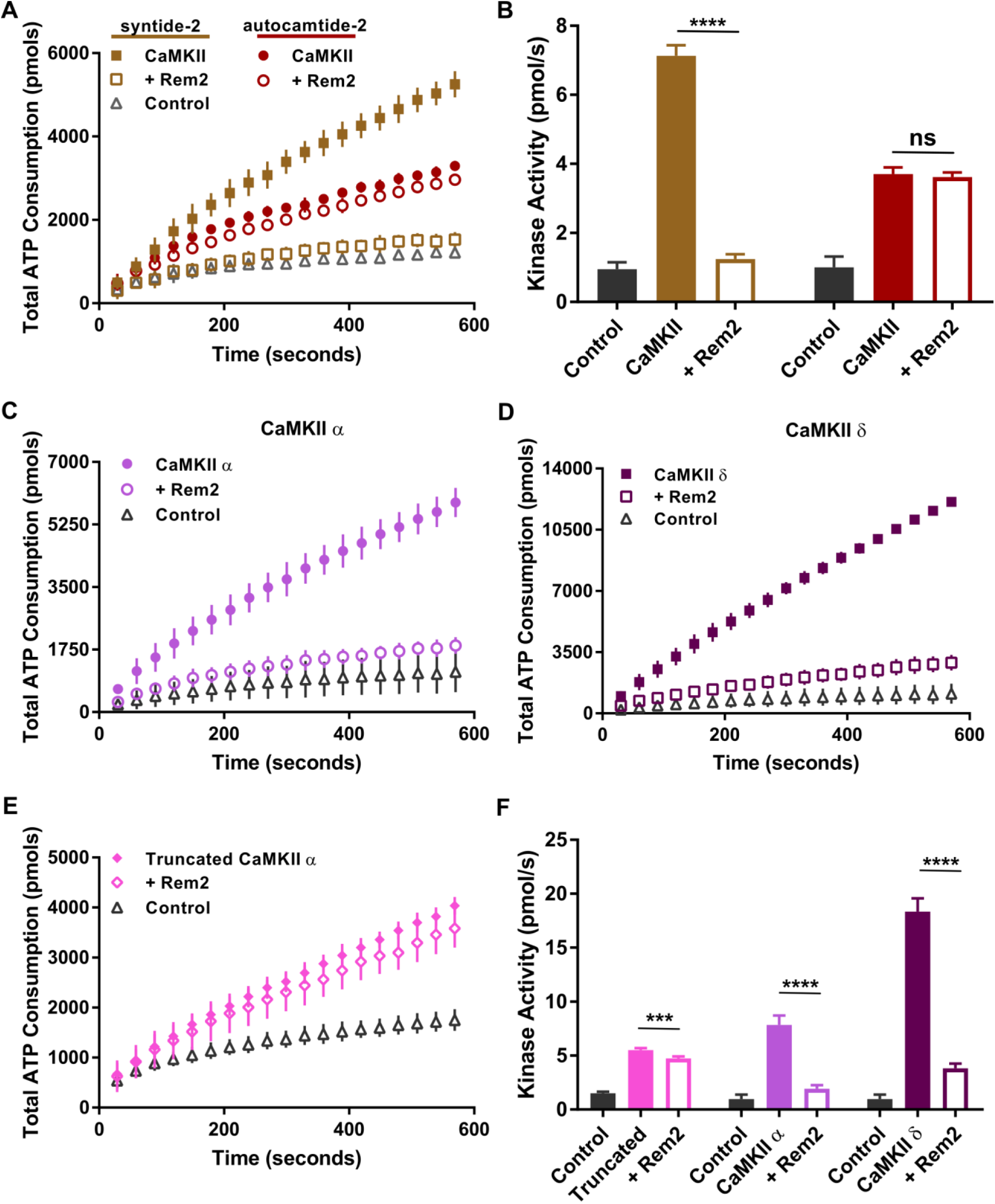
Conservation of Rem2 inhibition of CaMKII. (A) ATP consumption by purified CaMKIIα (20 nM monomer) monitored using the PK/LDH assay with syntide-2 (S-type substrate, 200 μM; gold filled squares) and autocamtide-2 (T-type substrate, 200 μM; red filled circles). Addition of 500 nM Rem2 produced a strong inhibition of syntide-2 phosphorylation (gold hollow squares) but not of autocamtide-2 (red hollow circles). Control is reaction minus CaMKIIα (gray triangles). For clarity, only the syntide-2 control is shown. (B) Quantification of the CaMKIIα kinase activity described in (A) using the same color code. (N = 5; standard deviations shown; ****, p< 0.0001; ns, not significantly different (p = 0.4449), using Student’s t-test). (C) and (D) ATP consumption by 20 nM purified CaMKIIα (Panel C, filled circles) and CaMKIIδ isoforms (Panel D, filled squares) onto syntide-2 (200 μM) in the absence or presence of Rem2 (500 nM, hollow circles and squares, respectively). Negative controls (gray triangles) did not contain the respective CaMKII isoform. (E) ATP consumption by monomer of rat CaMKIIα that lacks the association domain (residues 1-325) (20 nM monomer) monitored using the PK/LDH assay with syntide-2 (200 μM) in the absence (filled diamonds) or presence of 500 nM Rem2 (hollow diamonds) (F) Quantification of the kinase activity of CaMKII shown in panels (C), (D), and (E) using the same color code (N = 6 for all groups; standard deviations shown; ***, p = 0.001; ****, p < 0.0001, using Student’s t-test).

We sought to gain insight into the functional implications of Rem2 inhibition of CaMKII by investigating whether Rem2 inhibition is CaMKII isozyme specific, or if Rem2 is able to inhibit multiple CaMKII isozymes. Rem2 is expressed in several tissues including the brain, where all four CaMKII isozymes are present (47). Interestingly, peptides from all four CaMKII isozymes were identified in our mass spectrometry analysis, suggesting that Rem2 does not interact exclusively with the α isoform. Thus, we chose to extend our inhibition studies to the CaMKIIδ isozyme, which is expressed in neuronal and non-neuronal cells, but is also the major cardiac CaMKII (47) (Fig. 3C-D, F, Fig. S4B). We found that Rem2 is also a potent inhibitor of CaMKIIδ, similar to its effect on CaMKIIα, demonstrating that Rem2 inhibition of CaMKII is not restricted to the α isoform. This finding raises the possibility that inhibition of CaMKII might be a conserved feature of other RGK family members.

As CaMKIIα and CaMKIIδ differ by only a few amino acids in the variable domain, the inhibition data argue that the regions required for Rem2 inhibition are likely contained within conserved segments of the protein. To test this idea we used a monomeric truncated version of CaMKII that contains only the catalytic domain and Ca^2+^/CaM binding region. Rem2 displays only a minor inhibitory effect on syntide-2 phosphorylation using this truncated version of CaMKIIα (Fig. 3E-F, Fig. S4B). This result strongly suggest that the inhibition requires the association domain and is specific for the holoenzyme, the physiologically relevant form of the enzyme.

In this report, we demonstrate that Rem2 is a potent, endogenous inhibitor of CaMKII activity. These data represent a paradigm shift in our understanding of both RGK family and CaMKII function. Interestingly, Flynn *et al*. (2012) observed an NMDAR-dependent clustering of Rem2 and CaMKII in neurons, suggesting that the association of these proteins is responsive to changes in cellular activity. Thus, one possibility is that upon neuronal depolarization, Rem2 acts in a negative feedback loop to control autonomous CaMKII activity, perhaps modulating long-term potentiation or other cellular outputs of CaMKII signaling.

To date, only the 79 amino acid CaMKIIN isoforms α and β have been demonstrated to be endogenous inhibitors of CaMKII in mammals (25,26). However, the physiological role of CaMKIIN and the biological significance of the CaMKII-CaMKIIN interaction, remain unclear. Work from our lab using loss of function approaches in neurons demonstrated that Rem2 is a positive regulator of synapse formation and dendritic branching (36,37). Going forward these findings provide a framework for understanding the physiological significance of Rem2 inhibition of CaMKII in the context of synapse formation and dendritic branching.

Furthermore, our findings may provide a deeper understanding of calcium channel regulation. The inhibition of high-voltage Ca^2+^ channels by overexpression of Rem2 and other members of the RGK family have been extensively characterized. However, neither the precise mechanism of inhibition is known, nor is the relevance of this inhibition understood in more physiological context. CaMKII is an important modulator of high-voltage calcium channels and is directly involved in the frequency facilitation of the channel: phosphorylation of channels by CaMKII affects the channel conductance and inactivation properties (48-55). We speculate that by being a powerful inhibitor of both, Rem2 might facilitate the interaction between CaMKII and high-voltage calcium channels, perhaps by acting as a scaffold to bridge between CaMKII and VGCCs, thereby providing a layer of channel regulation that is activity-dependent.

Interestingly, the RGK family member Rad interacts with CaMKIIδ in cardiac myocytes and further, Rad knockdown leads to increased CaMKII activity in these cells (56,57). Thus, it is possible that inhibition of CaMKII is a conserved function of RGK family members and is relevant to other tissues where the RGK family members are strongly expressed: heart muscle, skeletal muscle, kidney, and pancreas. In particular, we hypothesize that understanding RGK regulation of CaMKII in other excitable cell types such as heart muscle, where Ca^2+^ signaling plays a critical role in the functional output of these cells, will yield important insights into the physiology of these cells.

## Materials and Methods

Detailed Materials and Methods are described in the Supplemental Data.

### CaMKII Activity Assays

The kinase activity of CaMKII was assessed using a continuous spectrophotometric assay. The standard assay (150 μL total volume) contained: 50 mM Tris (pH 7.5), 150 mM NaCl, 2 mM MgCl_2_, 1 mM CaCl_2_, 3.33 μM CaM, 400 μM sodium ATP, 200 μM phosphoenolpyruvate, 400 μM NADH, 9-15 units of pyruvate kinase, 13.5-21 units of lactate dehydrogenase, and 40 μM of either GDP-β-S (Fig. 1C, 1 G, 2A-E, 3A-E) or GMP-PCP (Fig. 1D). The substrate peptides used were either syntide-2 (PLARTLSVAGLPGKK) or autocamtide-2 (AC-2 - KKALRRQETVDAL), both at 200 μM. The reaction was started by the addition of 20 nM CaMKII (monomer concentration) and the decrease in absorbance at 340 nm at room temperature monitored in a reaction was stopped by addition human CaMKIIα and δ were purified as described in the supplemental data. Rat truncated CaMKIIα was obtained from New England Biolabs (Ipswich,MA). A calibration curve was used to convert the NADH absorbance readings at 340 nm into moles of ATP based on the 1:1 coupling of NADH:ATP consumption of the PK/LDH assay. Subsequently, the moles of ATP consumed in each step were used to construct a cumulative plot of ATP consumption. Then the kinase activity in pmols/s was calculated from the slopes of this plot.

### Autonomous CaMKII

In order to obtain autonomous CaMKII activity (i.e. Ca^2+^-independent kinase activity), the purified enzyme (60 nM) was pre-incubated with sodium ATP (500 μM) and CaM (10 μM) in 50 mM Tris buffer (pH 7.5) containing 150 mM NaCl, 1 mM CaCl_2_, and 2 mM MgCl_2_ in a total volume of 50 μL. After 2 min, 6 μL of 100 mM sodium EGTA was added to stop the autophosphorylation of CaMKII. The reaction mix was immediately added to 96 μL of a solution containing 200 μM syntide2, 400 μM NADH, 200 μM phosphoenolpyruvate, 500 μM sodium ATP, 40 μM GDP-β-S, 9-15 units of pyruvate kinase, and 13.5-21 units of lactate dehydrogenase in 50 mM Tris buffer (pH 7.5) containing 150 mM NaCl. Addition of sodium EGTA lowered the final concentration of free Ca^2+^ to < 20 nM while decreasing the free Mg^2+^ to ~ 1.2 mM.

### Radioactive in vitro Kinase Assay

The kinase activity of CaMKIIα was monitored by measurement of the incorporation of radiolabeled phosphate from [γ-^32^P]ATP into the substrate peptide syntide-2. The kinase assay (150 μL total volume) contained 50 mM Tris (pH 7.5), 150 mM NaCl, 2 mM MgCl_2_, 1 mM CaCl_2_, 3.33 μM CaM, 400 μM [γ-^32^P]ATP (~0.8 Ci/mmol), 200 μM syntide-2 peptide, and 40 μM of either GDP-β-S or GMP-PCP. The reaction was initiated by addition of purified CaMKII (20 nM final) and incubated at room temperature for 8 min. The reaction was stopped by addition of 600 μL 75 mM phosphoric acid and slot-blotted onto Whatman P81 phosphocellulose paper. After additional washes with 75 mM phosphoric acid, the paper was rinsed in acetone, dried and the radioactivity of bound syntide-2 measured using a phosphorimager. The intensities of the detected bands were quantified using the software ImageJ (NIH, Bethesda, MD).

### CaMKII Autophosphorylation

CaMKII autophosphorylation was assessed by measurement of the incorporation of radiolabeled phosphate from [γ-^32^P]ATP. The assay contained 50 mM Tris (pH 7.5), 150 mM NaCl, 2 mM MgCl_2_, 1 mM CaCl_2_, 3.33 μM CaM, 100 μM [γ-^32^P]ATP (~16 Ci/mmol), and 40 μM GMP-PCP. The reaction was initiated by addition of CaMKII (80 nM final) and incubated at 30^o^C for 2 min. The reaction was stopped by addition of 4x Laemmli loading buffer followed by incubation at 90 °C for 2 min. The samples were separated by SDS-PAGE and the gel blotted onto a Whatman 1 filter paper and dried. Radioactivity of protein bands was measured using a phosphorimager. The intensities of the detected bands were quantified using the software ImageJ (NIH, Bethesda, MD).

## Acknowledgements

We thank Dr. Leslie C. Griffith for critical comments on the manuscript. We are grateful to Dr. Stephen Van Hooser and members of the Paradis and Marr labs for resources and discussions. This work was supported by NIH grants R01NS065856 (S.P.), R21GM117034 (M.T.M), and R01GM117034 (M.T.M.). The content is solely the responsibility of the authors and does not necessarily represent the official views of the National Institutes of Health.

## Author Contributions

L.R, J.J.H., M.T.M., and S.P. designed research; L.R, J.J.H., K.K., B.T., J.C.C., and M.T.M. performed research; L.R, J.J.H., and M.T.M. analyzed data; L.R., M.T.M., and S.P. wrote the paper.

## Footnotes

The abbreviations used are: CaM, calmodulin; CaMKII, Ca2+/calmodulin-dependent protein kinase II; CREB, cAMP response element-binding protein; GAP, GTPase-activating protein; GDP-β-S, Guanosine 5^′^-[β-thio]diphosphate; GEF, guanine nucleotide exchange factor; GMP-PCP, Guanosine-5’-[(β,γ)- methyleno]triphosphate; LTP, long-term potentiation; MBP, maltose-binding protein; PK/LDH, pyruvate kinase/lactate dehydrogenase.

